# Mutant ACTB mRNA 3′UTR Promotes Hepatocellular Carcinoma Development by Regulating miR-1 and miR-29a

**DOI:** 10.1101/691527

**Authors:** Yong Li, Hong-Bin Ma, Chang-Ying Shi, Fei-Ling Feng, Liang Yang

## Abstract

In recent years, mounting studies have shown that ACTB is closely related to various tumors. Although ACTB is dysregulated in numerous cancer types, limited data are available on the potential function and mechanism of ACTB in hepatocellular carcinoma (HCC). This study evaluated the expression and biological roles of mutant ACTB mRNA 3′UTR in HCC. Transcriptome sequence and qRT-PCR analysis determined that mutant ACTB mRNA 3′UTR was high expression in HCC tissues. Luciferase reporter assay showed that the ACTB mRNA 3′UTR mutations made it easier to interact with miR-1 and miR-29a. Moreover, mutant ACTB mRNA 3′UTR regulated miR-1 and miR-29a degradation via AGO2. Furthermore, mutant ACTB mRNA 3′UTR promoted hepatocellular carcinoma cells migration and invasion *in vitro* and *in vivo* by up-regulating miR-1 target gene MET and miR-29a target gene MCL1. In a word, our study demonstrates that 3′UTR of ACTB plays a key role in the tumor growth of hepatocellular carcinoma (HCC) and highlights the molecular mechanisms of ACTB-involved cancer growth and development.

## Introduction

MicroRNAs (miRNAs) are the key regulators of genes expression by binding to the 3’-untranslated region (UTR) of the target mRNA through seed-match sequences[1, 2], leading to the target mRNA translational repression or cleavage[3]. The 3′UTR single nucleotide polymorphisms in the mRNA-miRNA seed-match regions, which is shorten for miRNA related SNP can impact target genes expression[4], causing signaling pathways changes[5]. Since some of these signals may be associated with tumorigenesis[6, 7], polymorphisms in the miRNA-binding site of the target gene can eventually affect individual’s cancer risk and perhaps can be used for prognostic prediction[8].

Housekeeping genes which are necessary for basic cell survival and present in all nucleated cells are commonly used as endogenous references. We take it for granted that housekeeping genes are not affected by any human diseases. However, numerous studies have shown that many of the commonly used housekeeping genes are regulated and vary under different physiological and pathological conditions[9–11].

Beta-actin (ACTB) has been regarded as an endogenous housekeeping gene all the time and has been widely used as a reference gene/protein in quantifying expression levels in cells or tissues. However, accumulating evidence indicates that ACTB is closely associated with the development of multiple cancers[12, 13], such as liver carcinoma, melanoma, colorectal cancer and so on. Several studies have shown a correlation of ACTB with liver cancer development and metastasis[14]. ACTB was de-regulated in different TNM stages and degrees of tumor invasiveness of HCC[15, 16]. However, the detailed mechanism of ACTB in liver carcinoma progression is still unclear and worthy of study.

In this study, we show that mutant ACTB mRNA 3′UTR promotes the tumor growth of hepatocellular carcinoma by regulating miR-1 and miR-29a degradation via AGO2, thus up-regulating miR-1 target gene MET and miR-29a target gene MCL1 expression. These findings provide a new insight on molecular mechanisms of ACTB-involved cancer growth and development.

## Results

### The mRNA level of housekeeping gene ACTB is elevated in hepatocellular carcinoma along with high frequency mutation of 3′UTR

It has been reported that housekeeping genes are involved in different physiological and pathological conditions, for example, ACTB is de-regulated in hepatocellular carcinoma. We performed transcriptome sequence to analyze the expression of housekeeping genes in 88 pairs of human HCC specimens. The top 9 housekeeping genes with high UTR mutation frequency was listed in Figure 1, in which ACTB listed first for the reason that the same mutation site exists in 21 pairs of samples. We further analyzed the mutation of ACTB 3′UTR in details, as the DNA mutation sites and corresponding mRNA mutation sites shown in Table 1. The two sites with the maximum mutation frequency were 5527500 with A inserted and 5527291 C>G, correspondingly, the mutation at mRNA was A inserted (1567-1568) and 1777 C>G.

**Figure 1.**
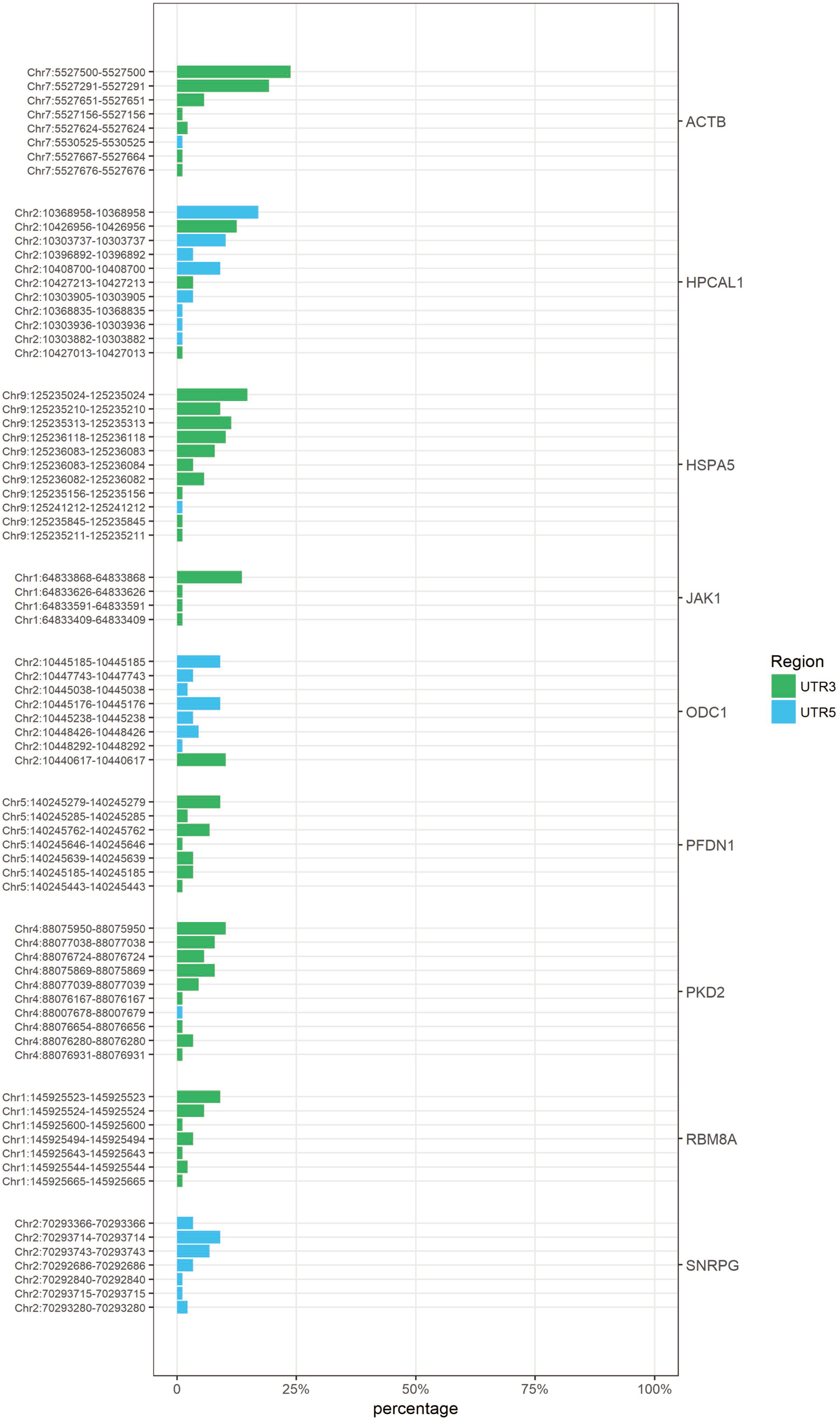
Identification of ACTB mRNA 3′UTR mutant is the most often among housekeeping genes mutation in hepatocellular carcinoma. The mutation sites are on the left. The mutant housekeeping genes are on the right. The data are listed the top 9 housekeeping genes in descending order of mutation frequency analyzed by transcriptome sequence.

**Table 1.**
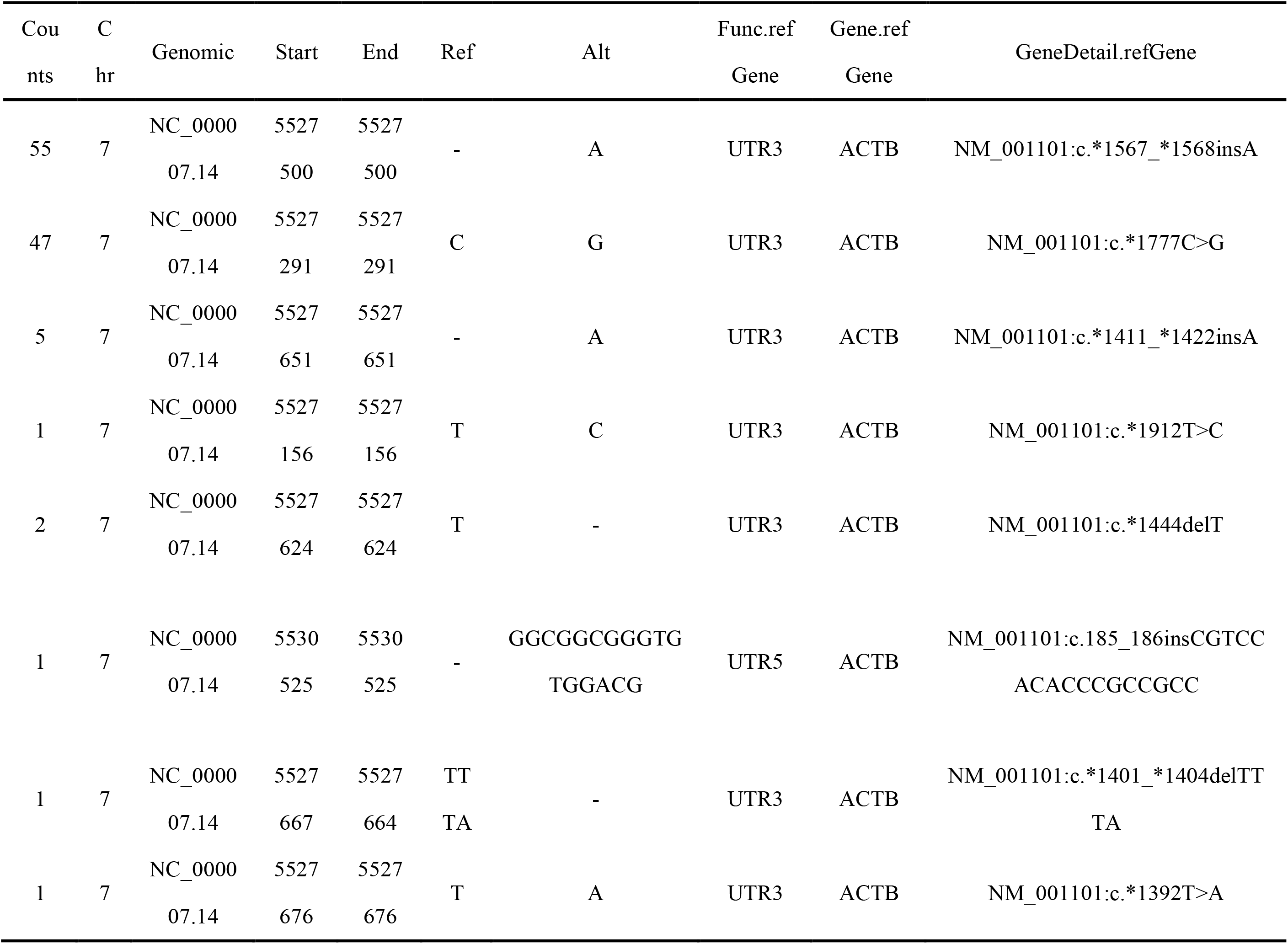
DNA mutation sites and corresponding mRNA mutation sites of ACTB 3’-UTR in hepatocellular carcinoma.

Then we examined ACTB expression in hepatocellular carcinoma (HCC) tissues. The copy number variation of ACTB was analyzed using Oncomine. The result showed that there was no significant difference in copy number variation of ACTB between tumor tissues (n = 97) and normal control (n = 115) (Fig 2A). Meanwhile, the relative expression of ACTB at mRNA level was increased in HCC tissues (n=226) than normal control (n=220) (Fig 2B). Next, we verified ACTB mRNA expression in tissue using qPCR. To ensure the data reliability, we used 18S and GUSB as internal reference, respectively. Consistent with the bioinformatics analysis through Oncomine, the relative mRNA expression of ACTB in tumor tissues among 20 cases was significantly higher than that of corresponding adjacent non-tumor tissues (Fig 2C and 2D). Interestingly, although the ACTB mRNA level was increased, there was no obvious difference between tumor tissues and the corresponding normal tissues at protein level (Fig 2E)

**Figure 2.**
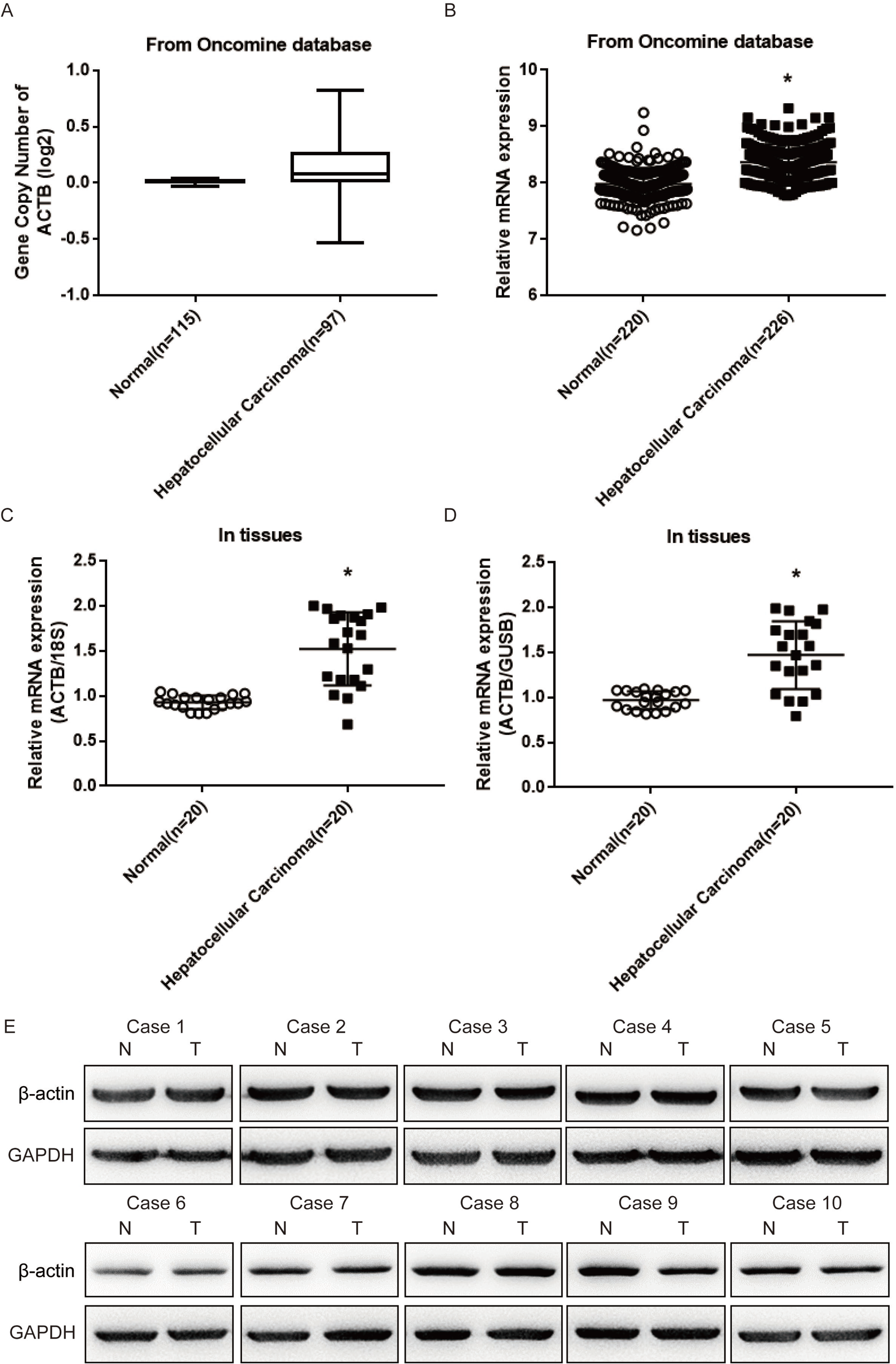
ACTB is overexpressed at mRNA level in hepatocellular carcinoma tumor tissues with no obvious change at copy number variation (CNV). A, B ACTB copy number variation in HCC tissues is similar to the non-tumor tissues (A) whereas ACTB mRNA level is significantly higher in HCC compared with paired non-tumor tissues in a data set from the Oncomine database (B), \**P* <0.05. C, D ACTB mRNA levels are detected by qRT-PCR quantified by endogenous control 18S (C) or GUSB (D) in randomly selected HCC and paired non-tumor tissues. \**P* <0.05. E β-actin protein level in randomly selected HCC and paired non-tumor tissues is detected by western blot.

### ACTB mRNA 3′UTR mutation affects the interaction with miRNA

To study the role of mutant ACTB mRNA 3′UTR in the hepatocarcinoma development, we focused on the microRNA which might bind to the target sites of 3′UTR. We found that the inserted mutation of 3′UTR at 1567 was miR-1 potential binding sites, resulting in miR-1-3p base pairing with the region from 7 bp to 8 bp. Another mutation led to a potential binding site for miR-29a-3p (Fig 3A). It was analyzed that the total free energy of potential binding sites interaction between miR-1 and miR-29a with ACTB mRNA 3′UTR using RNAhybrid (https://bibiserv.cebitec.uni-bielefeld.de/rnahybrid). The results showed that the ACTB mRNA 3′UTR mutation reduced the free energy of interaction with microRNAs (Fig 3B), revealing that mutations might change the interaction between microRNAs and their target sites. To confirm that the mutation of ACTB mRNA 3′UTR changed the interaction with miR-1 and miR-29a, we constructed the luciferase reporter plasmids containing wild type / mutant ACTB mRNA 3’ UTR. The luciferase reporter assay showed that, compared with wild type 3′UTR, miR-1 and miR-29 significantly repressed the luciferase activity of which containing the mutant 3′UTR, as shown in Fig 3C, suggesting that the ACTB mRNA 3′UTR mutations made it easier to interact with miR-1 and miR-29a.

**Figure 3.**
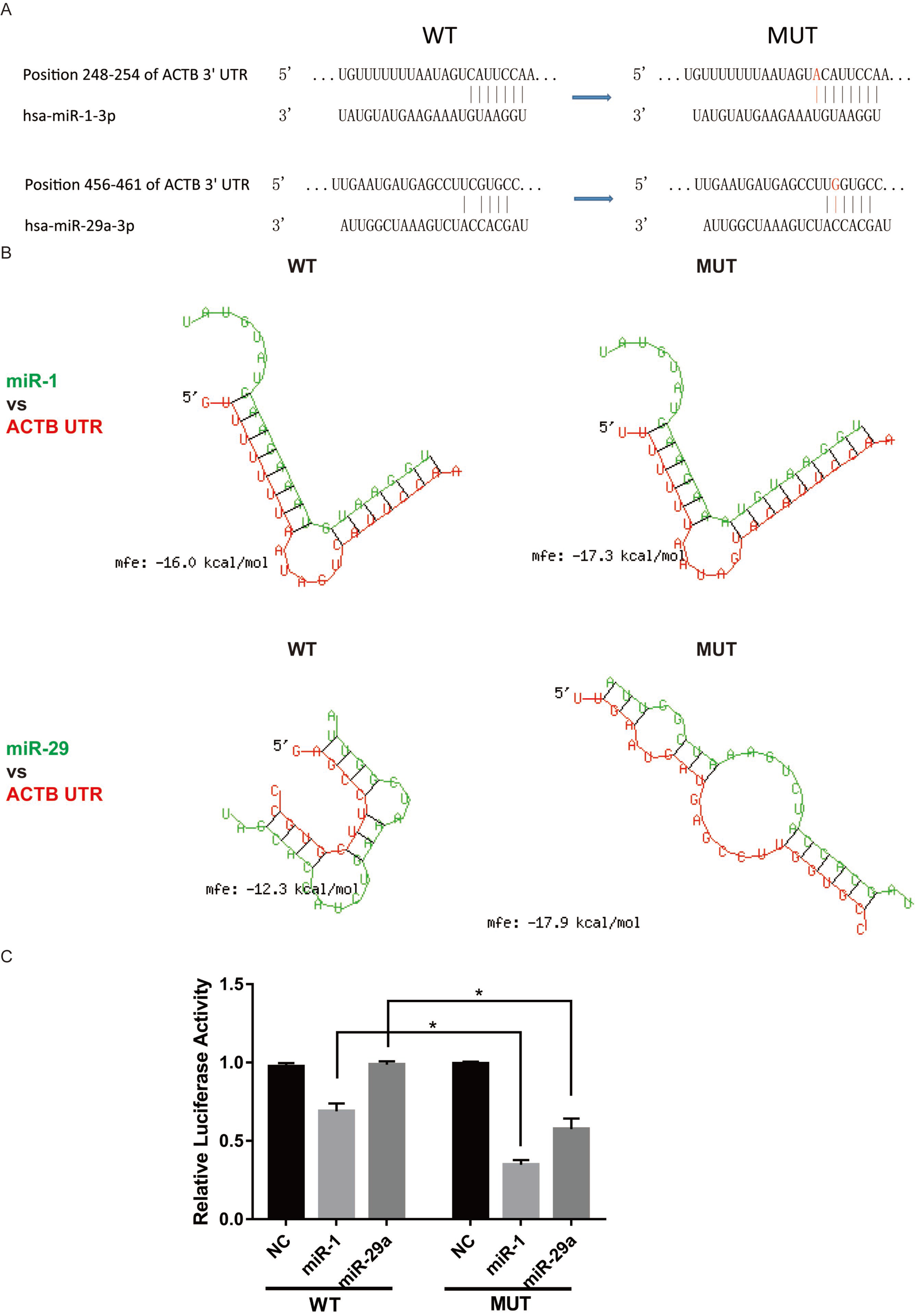
Mutant ACTB 3′UTR is liable to interact with miRNA. A Schematic of the SNP of the 3′UTR of ACTB (mutated nucleotides shown in red) promoting the interaction with miR-1 and miR-29a respectively, by complementary pairing with more predicted binding sites of miR-1 and miR-29a. B Schematic of the secondary structure formed between miR-1/ miR-29a and the ACTB 3′UTR wildtype or mutant and the free energy of interactions. C HepG2 cells were cotransfected with different miRNAs and the luciferase reporter constructs harboring ACTB or mutant ACTB 3′UTR fragments. A non-related fragment of RNA was used as a control. Luciferase activity assays indicated that miR-1 and miR-29a significantly repressed luciferase activities when the luciferase constructs harbored the mutant ACTB 3′UTR compared to ACTB. \**P*<0.05. All experiments were performed in triple.

### The mutant ACTB mRNA 3′UTR affects the relative expression of mature miR-1 and miR-29a

In order to investigate the effects made by ACTB mRNA 3′UTR mutation in hepatocellular carcinoma, we performed the qPCR to detect the relative expression of miR-1 and miR-29a in liver cancer tissues. It turned out that miR-1 and miR-29a were significantly decreased in the liver cancer tissues as shown in Figure 4A. Since miRNA destabilization through endogenous targets (endogenous target-RNA-directed miRNA degradation, TDMD) emerges as an effective post-transcriptional mechanism for the selective regulation of specific miRNAs[17, 18], we wondered if miRNA’s reduction is related to the mutation in ACTB mRNA 3′UTR. In human hepatocellular carcinoma cells Hep3B and HepG2, we examined the relative expression of miR-1 and miR-29a by transfection with plasmids containing wild-type or mutant ACTB mRNA 3′UTR, respectively. The results showed that mature miR-1 and miR-29a expression did decrease with mutant ACTB mRNA 3′UTR overexpression (Fig 4B and 4C), but neither pre-miRNA nor pri-miRNA levels changed (Fig 4D-G). These results suggested that the mutant ACTB mRNA 3′UTR can reduce the relative expression of mature miR-1 and miR-29a, but are not related to the transcriptional regulation and miRNA precursor processing of miR-1 and miR-29a.

**Figure 4.**
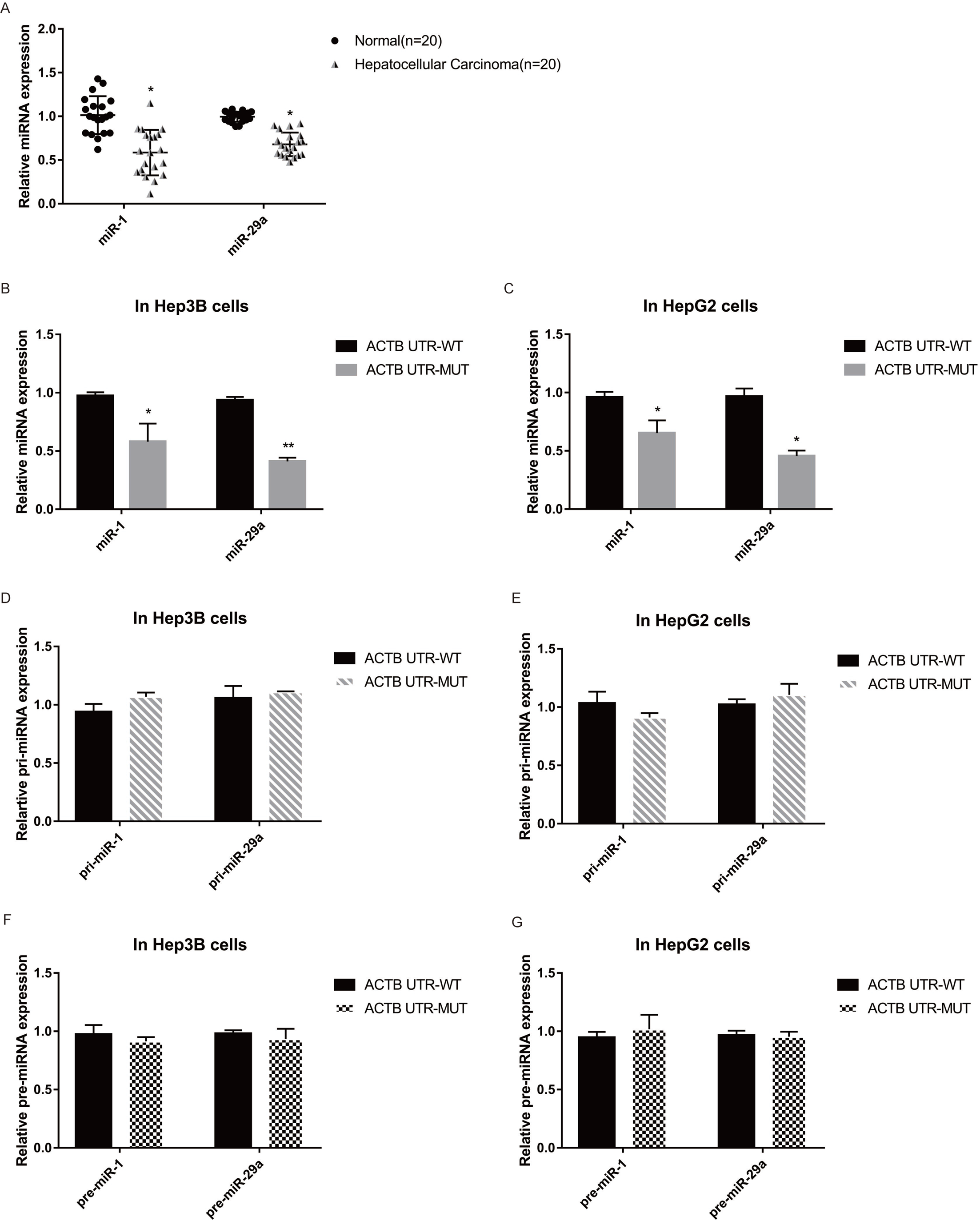
Mutant ACTB 3′UTR modulates the activity of miR-1 and miR-29a. A miR-1 and miR-29a are lower-expression in HCC tumor tissues compared with paired non-tumor tissues by qRT-PCR. \**P*<0.05. B, C The mRNA level of miR-1 and miR-29a was down-regulated by mutant ACTB 3 ′ UTR compared with wildtype control both in Hep3B (B) and HepG2 (C). All experiments were performed in triple. D, E The pri-RNA level of miR-1 and miR-29a has no difference between cells transfected with ACTB and mutant ACTB 3′ UTR both in Hep3B (D) and HepG2 (E). \**P*<0.05. All experiments were performed in triple. F, G The pre-RNA level of miR-1 and miR-29a shows no obvious change between cells transfected with ACTB and mutant ACTB 3′ UTR both in Hep3B (F) and HepG2 (G). \**P*<0.05. All experiments were performed in triple.

### AGO2 is involved in the mutant ACTB mRNA 3′UTR mediated miRNA degradation

To further explore the mechanism that the mutant ACTB mRNA 3′UTR affected the relative expression of mature miR-1 and miR-29a, we conducted miRNA stability tests. We found that the stability of miR-1 and miR-29a significantly decreased in cells that expressed the mutated ACTB mRNA 3′UTR after the inhibition of intracellular RNA transcription with prolonged actinomycin D treatment (Fig 5A and 5B). These results suggest that the mutant ACTB mRNA 3′UTR may induce target-directed miRNA degradation (TDMD), which is a post-transcriptional regulation of miR-1 and miR-29a. However, it is still not clear about the regulatory mechanism of TDMD. Haas et al. found that exonuclease DIS3L2 was involved in TDMD through interaction with AGO2. We used siRNA to silence the expressions of DIS3L2 and AGO2 respectively (Fig 5C and 5D), and the results showed that reducing the expression of DIS3L2 did not inhibit the degradation of miR-1 and miR-29a induced by mutant ACTB mRNA 3′UTR (Fig 5E and 5F). However, AGO2 down-regulation can partially restore the stability of miR-1 and miR-29a (Fig 5E and 5F). Furthermore, we analyzed the relative expression of DIS3L2 and AGO2 at mRNA and protein levels in liver cancer tissues (Fig 5G-I), and the results showed that AGO2 was highly expressed in liver cancer tissues compared to normal tissues at both mRNA and protein levels. These results indicated that AGO2 was involved in the TDMD of which mutant ACTB mRNA 3’-UTR on miR-1 and miR-29a.

**Figure 5.**
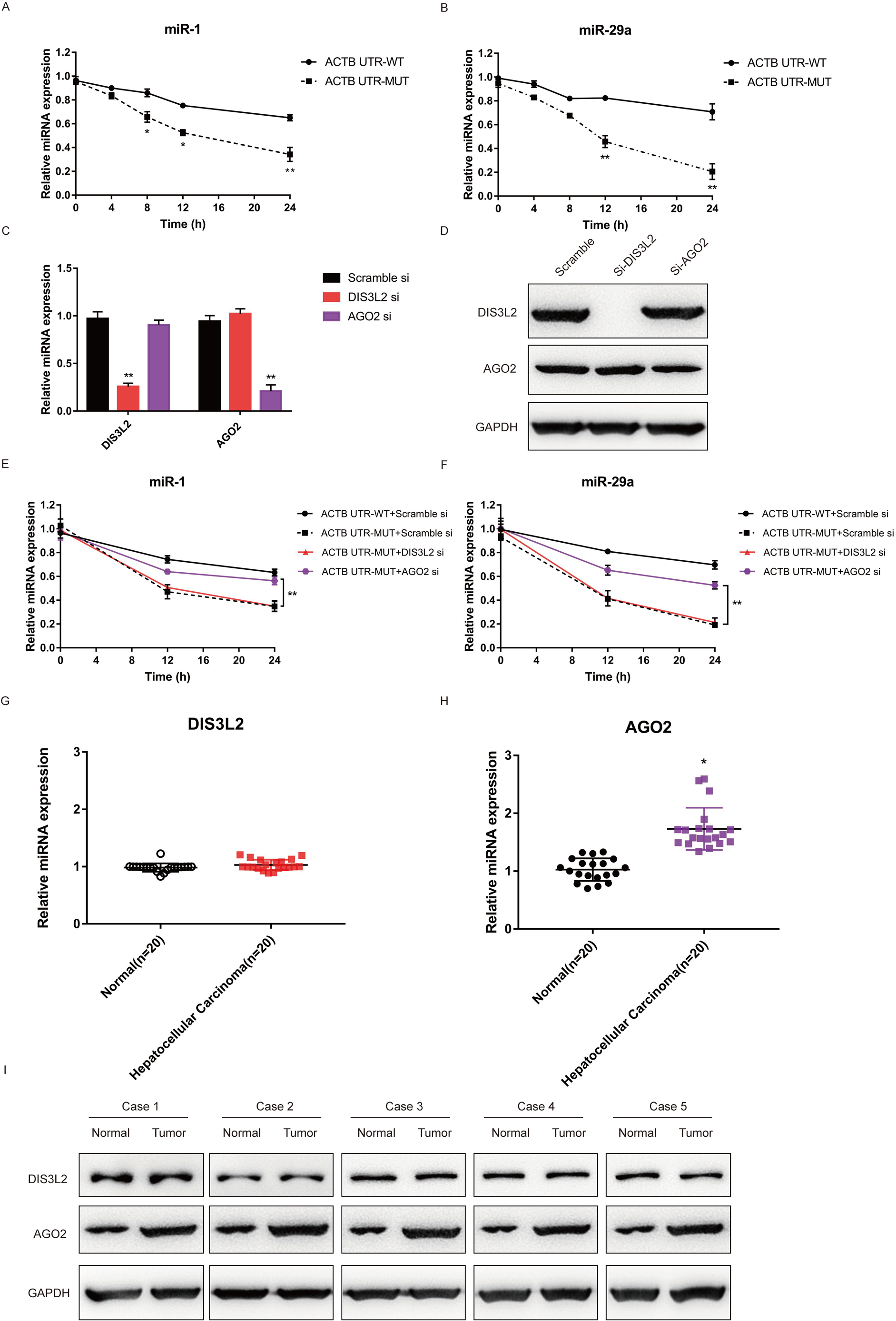
Mutant ACTB 3′ UTR regulates miRNA stability via AGO2. A, B The expression of miR-1 (A) and miR-29a (B) is decreased even more pronounced with the increased transfection time of mutant ACTB 3′ UTR. \**P*<0.05, \*\**P*<0.01. C The knockdown efficiency of siDIS3L2 or siAGO2 at mRNA level is detected by qRT-PCR. \**P*<0.05, \*\**P*<0.01. D The knockdown efficiency of siDIS3L2 or siAGO2 at protein level is detected by western blot. E, F The decreased expression of miR-1 (E) and miR-29a (F) is partially rescued by loss of AGO2 but not DIS3L2. \**P*<0.05, \*\**P*<0.01. G, H The expression of DIS3L2 (G) and AGO2 (H) is examined by qRT-PCR in randomly selected HCC and paired non-tumor tissues. \**P*<0.05. I Expression of DIS3L2 and AGO2 at protein level is examined by western blot in randomly selected HCC and paired non-tumor tissues. All experiments were performed in triple.

### The mutant ACTB mRNA 3’-UTR mediates the cellular biological function of HCC

To further explore the functional effects induced by the mutated ACTB mRNA 3’-UTR through mediating miR-1 and miR-29a TDMD, we overexpressed the mutant ACTB mRNA 3′UTR in human hepatocarcinoma cells. We observed that the protein levels of both miR-1 potential target MET and miR-29a potential target MCL1 were significantly increased in Hep3B[19, 20] (Figure 6A), accompanied by enhanced cell proliferation (Fig 6B and 6C). In addition, the migrating and invasive ability of mutant ACTB mRNA 3′UTR-overexpressed cells was also significantly enhanced compared to control (Fig 6D-G). These results suggested that the mutated ACTB mRNA 3′UTR promoted the proliferation, migration and invasion of hepatocellular carcinoma cells by mediating the TDMD effect of miR-1 and miR-29a, leading to the up-regulation of target genes of miR-1 and miR-29a.

**Figure 6.**
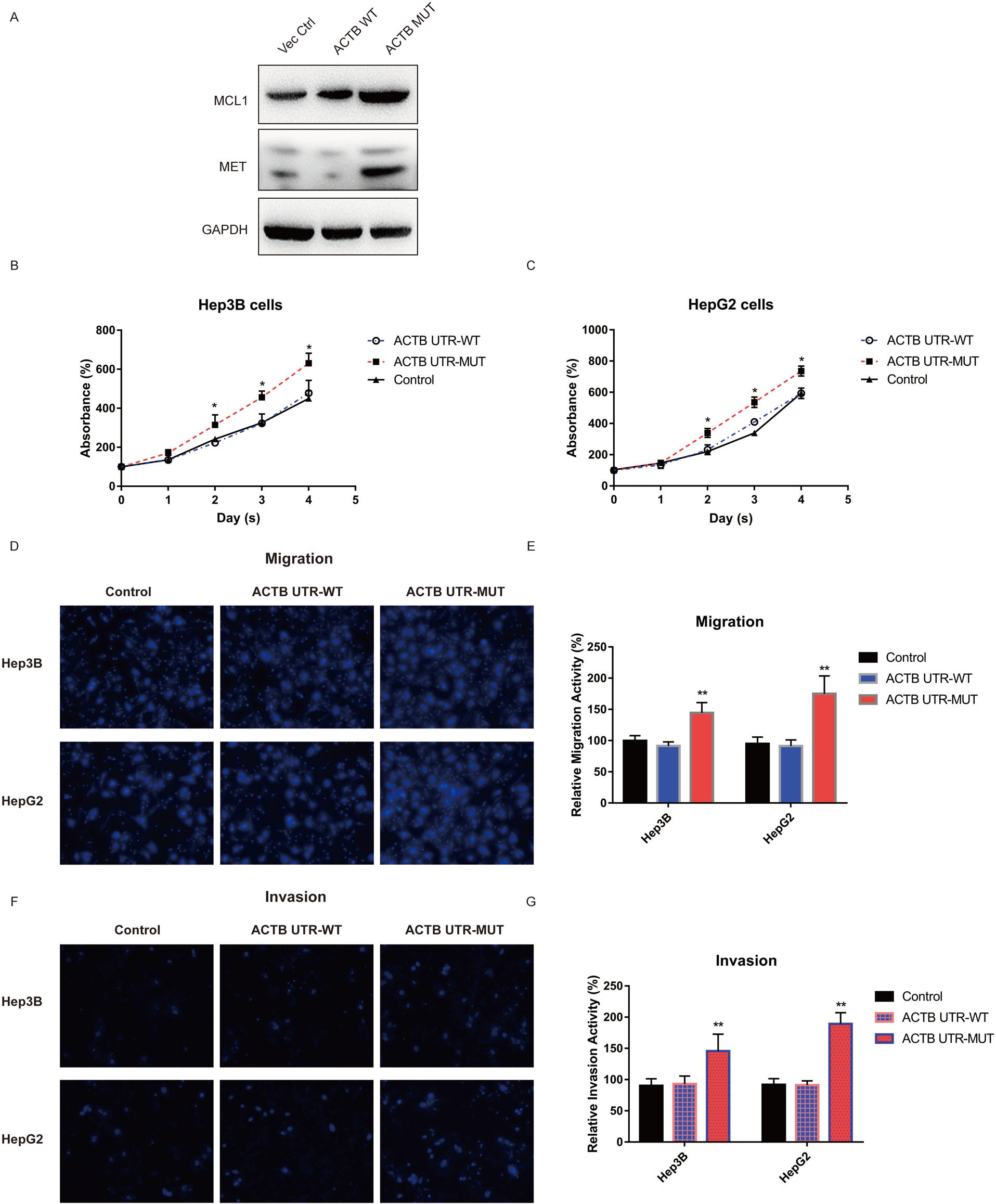
Mutant ACTB 3′ UTR promotes the malignant phenotype of hepatocellular carcinoma cells. A The protein expression of MCL1 and MET is detected by western blot after cells transfected with ACTB 3′ UTR wildtype and mutant in Hep3B cells. B, C The cell proliferation is analyzed by CCK8 after transfection with ACTB, mutant ACTB 3′ UTR or vector control in Hep 3B (B) and Hep G2 (C). Mutant ACTB 3′ UTR promotes hepatocellular carcinoma cells proliferation within 5 days. D-G Cells were transfected with ACTB, mutant ACTB 3′UTR or vector control, respectively. The influence of mutant ACTB 3′ UTR on the migration (D, E) and invasion (F, G) of hepatocellular carcinoma cells was analyzed by transwell migration and invasion assays, respectively. ** *p* <0.01. Mutant ACTB 3′ UTR significantly promotes hepatocellular carcinoma cells migration and invasion. All experiments were performed in triple.

### ACTB mRNA 3′UTR mutant promotes hepatocellular carcinoma tumorigenesis *in vivo*

To determine the role of the mutated ACTB mRNA 3′UTR in tumor development *in vivo*, we used a xenograft mouse model. We constructed the luciferase lentivirus vector containing the wildtype or mutant ACTB mRNA 3′UTR. Hep3B cells stably transfected with wildtype ACTB or mutant ACTB mRNA 3′UTR were subcutaneously injected into male nude mice. We observed that the bioluminescence intensity of the mutant ACTB mRNA 3′UTR group was significantly higher than wildtype or controls in the Hep3B-derived xenografts (Fig 7). In conclusion, the mutant ACTB mRNA 3′UTR promotes tumor growth.

**Figure 7.**
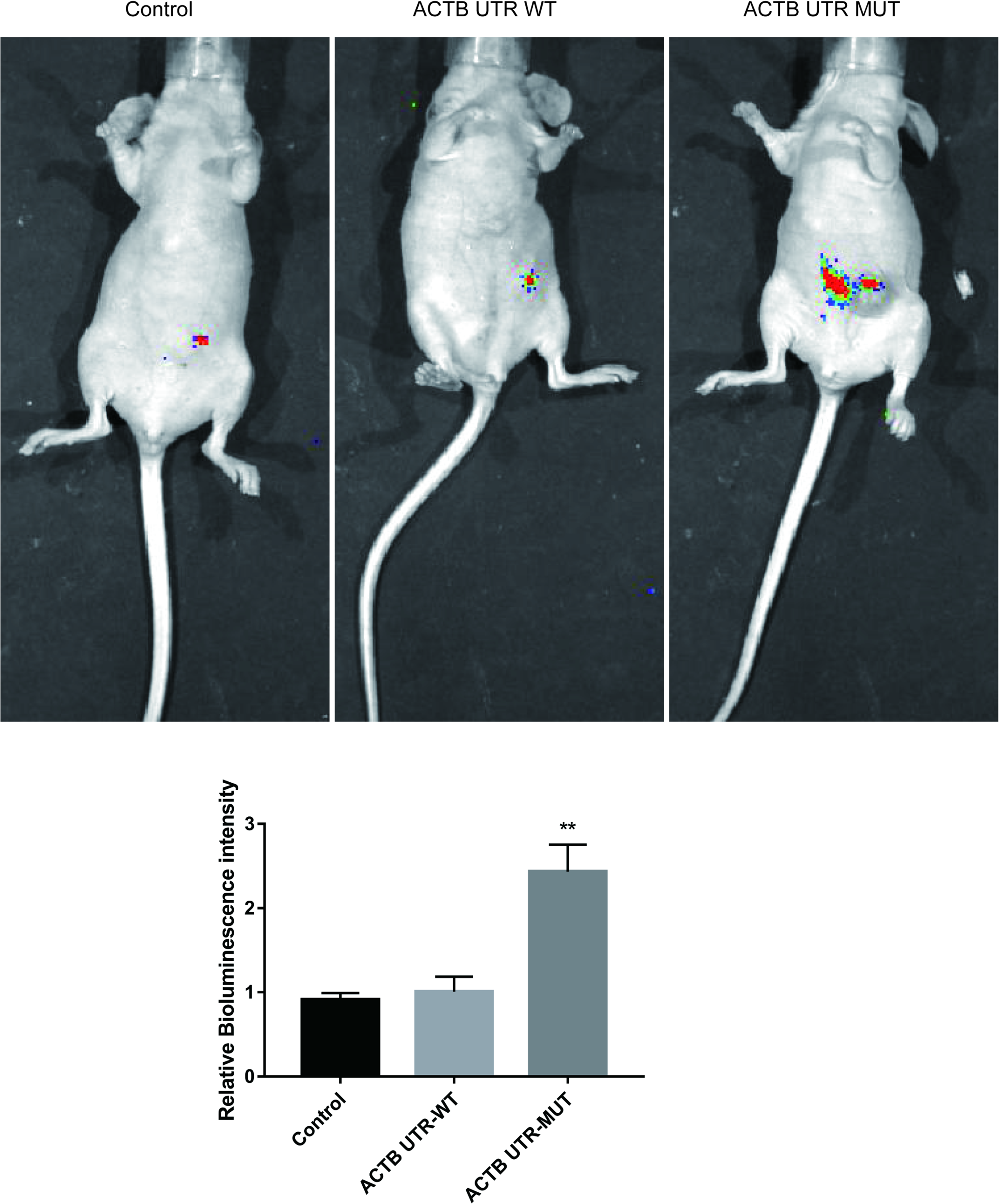
Mutant ACTB 3′ UTR promotes hepatocellular carcinoma tumorigenesis *in vivo*. Effects of mutant ACTB 3′UTR on localization of Hep3B cells in the liver of nude mice (n =5) at 8 weeks of injection of cells via intraperitoneal injection. The left panel shows bioluminescence imaging, and the right panel shows quantitative fluorescent intensities.

## Discussion

Over the past few decades, ACTB is commonly regarded as a constitutive housekeeping gene under the general circumstance, of which the expression is stable and unaffected by most physiological conditions. Therefore ACTB was widely used as an endogenous reference for genes quantification in multiple cells and tissues. However, mounting studies have shown that ACTB was regulated and vary under different experimental conditions, ACTB might be not suitable for gene level normalization[13]. For instance, ACTB mRNA levels in invasive melanoma cell T1C3 were two fold higher than that in non-invasive melanoma cell 1C8 melanoma cells derived from the same melanoma patient[21], implicating that ACTB was not a suitable reference gene/protein for melanoma cells from either the same patient or from heterogeneous cellular subpopulations of the same pathological origin. Rubie C et al. found that ACTB was up-regulated in pancreatic cancer (PC) at both RNA and protein levels, suggesting that ACTB was involved in PC and unsuitable as a reference gene/protein[9]. Several studies have shown that ACTB was up-regulated in HCV-induced and HBV-induced hepatocellular carcinoma (HCC), and overexpression of ACTB in tumorous tissues of HCC was with higher invasiveness and metastasis, indicating that ACTB might be closely associated with liver cancer growth and metastasis[14, 22, 23]. However, since ACTB are associated with hepatocellular carcinoma tumorigenesis, the function and regulatory mechanisms of ACTB in HCC progression remained to be explored. In the present study, we identified that the mutant ATCB mRNA 3′ UTR played a key role in the tumor growth in HCC by mediating miR-1 and miR-29a degradation.

The miR-1 in our study was miR-1-1. In humans there are two distinct microRNAs that share an identical mature sequence, which are called miR-1-1 and miR-1-2. We found that the mature miR-1-2 expression regulated by mutant ATCB mRNA 3′ UTR presented similar effects as miR-1-1 (Sup. Fig 1A), but with no altered transcriptional regulation and miRNA precursor processing (Sup. Fig 1B-C). Since miR-29b and miR-29c has the same seed region with miR-29a, we speculated that the stability of miR-29b and miR-29c might also be regulated by mutant ATCB mRNA 3′ UTR. The assumption was confirmed by our experiment as shown in Appendix Fig S2A and S2B.

AGO2 is essential for miRNA processing machinery and has been reported to be dysregulated in many types of cancers[24–27]. Wang M. et al. found that loss of miR-100 could enhance prostate cancer cells migration, invasion, epithelial-mesenchymal transition through targeting AGO2[28]. Völler D et al. revealed a strong reduction of AGO2 expression in melanoma compared with primary melanocyte[27]. Guo JF et al. reported that up-regulation of AGO2 contributed to cervical cancer malignancy[29]. In this study, we found that mutating the 3′ UTR of ACTB abolished miR-1 and miR-29a activity through AGO2. Silence of AGO2 could partially inhibit the degradation of miR-1 and miR-29a mediated by the mutant 3′ UTR.

Collectively, in our study, we identified that mutant ATCB mRNA 3 ′ UTR promoted hepatocellular carcinoma proliferation and invasion by AGO2-involved miR-1 and miR-29a degradation, thus enhancing MEL1 and MET expression. Our findings highlights on the molecular mechanisms of ACTB-involved cancer growth and development.

Taken together, our results revealed that the mRNA level of ACTB with hypermutation in 3′ UTR was increased in HCC tumor tissues and its upregulation may be related to tumorigenesis. The capacity of mutant ACTB 3 ′ UTR to promote HCC cell proliferation and migration may be achieved partially through repressing miR-1 and miR-29a expression via the interaction with AGO2. Generally, our study provides a novel perspective for the mechanism by which mutant ACTB 3′ UTR regulates HCC cell growth and migration.

## Materials and Methods

### Oncomine analysis

The expression levels of ACTB copy number and mRNA in HCC were analyzed using Oncomine[30]. For this, we compared tumor specimens with normal tissues datasets. We selected p<0.01 as a threshold to reduce the false discovery. We analyzed the results for fold changes and p-values.

### Patients and tissue samples

A total of 88 HCC patients at the Shanghai Eastern Hepatobiliary Hospital (China) between 2017 and 2018 were enrolled in this study. The patient characteristics were shown in Appendix Table S1. This study was approved by the Research Ethics Committee of Shanghai Eastern Hepatobiliary Hospital (EHBHKY2017-01-106), and written informed consent was obtained from all participants.

### Transcriptome sequence

RNA of the human HCC specimens was extracted for RNA-Seq analysis. RNA degradation and contamination was monitored on 1% agarose gels. RNA purity was checked using the NanoPhotometer® spectrophotometer (IMPLEN, USA). RNA concentration was measured using Qubit® RNA Assay Kit in Qubit® 2.0 Flurometer (Life Technologies, USA). RNA integrity was assessed using the RNA Nano 6000 Assay Kit of the Bioanalyzer 2100 system (Agilent Technologies, USA). A total of 3 μg RNA per sample was used as input material for the RNA sample preparations. Sequencing libraries were generated using NEBNext® Ultra™ RNA Library Prep Kit for Illumina® (NEB, USA) following manufacturer’s recommendations. The clustering of the index-coded samples was performed on a cBot Cluster Generation System using TruSeq PE Cluster Kit v3-cBot-HS (Illumina, USA). After cluster generation, the library preparations were sequenced on an Illumina Hiseq platform and 125/150 bp paired-end reads were generated. Reference genome and gene model annotation files were downloaded from genome website directly. Index of the reference genome was built using Bowtie v2.2.9 and paired-end clean reads were aligned to the reference genome using TopHat v2.1.1. HTSeq v0.6.1 was used to count the read numbers mapped to each gene. The fragments per kilobase million value of each gene was calculated based on the length of the gene and read count mapped to this gene. Differential expression analysis of two conditions was performed using the DESseq package (1.18.0). The resulting P-values were adjusted using the Benjamini and Hochberg’s approach for controlling the false discovery rate. Genes with an adjusted P-value <0.05 found by DESeq were assigned as differentially expressed.

### Cell culture

Human HCC cell lines (Hep G2 and Hep 3B) were purchased from the Institute of Biochemistry and Cell Biology at the Chinese Academy of Sciences (Shanghai, China). The cells were cultured in Dulbecco’smodified Eagle’s medium (DMEM) supplemented with 10% fetal bovine serum (10% FBS), 100 U/ ml penicillin, and 100 mg/ml streptomycin in incubators at 37°C with 5% CO2.

### RNA extraction and qRT-PCR analyses

The total RNA was extracted from cultured cells or frozen tissues using the TRIzol reagent (Invitrogen, Grand Island, NY, USA) according to the manufacturer’s instructions. For qRT-PCR, the isolated RNA was reverse transcribed to cDNA using a PrimeScript RT reagent Kit (Takara, Dalian China). Real-time PCR analyses were performed with SYBR Green (Takara, Dalian China). The results were normalized to the expression of 18S (ID: 106632259) and/or GUSB (ID: 2990). The gene-specific primers (5’-3’) were as follows:

miR-1 (NC_000020.11): Forward, CGCAGTGGAATGTAAAGAAG, and Reverse, GGTCCAGTTTTTTTTTTTTTTTATACATAC; miR-29a (NC_000007.14): Forward, CGCAGTAGCACCATCTGA; and Reverse, TCCAGTTTTTTTTTTTTTTTAACCGA; pre-miR-1: Forward, GGAAACATACTTCTTTATATGCCCAT, and Reverse, GTCCAGTTTTTTTTTTTTTTTGAGATAC; pre-miR-29a: Forward, TCTTTTGGTGTTCAGAGTCAATATAATTTTCTAGCACCATCT, and Reverse, TCCAGTTTTTTTTTTTTTTTATAACCGA; pri-miR-1: Forward, CGGGGTCTTGGAACTGCAT, and Reverse, TCCCGGCCTGAGATACATAC; pri-miR-29a: Forward, CGACCTTCTGTGACCCCTTA, and Reverse, TCATGGTGCTCTTCCCCAATC; >hsa-miR-29a-3p MIMAT0000086: UAGCACCAUCUGAAAUCGGUUA; qRT-PCR and data collection were carried out on an ABI 7500 real-time PCR system (Applied Biosystems, Foster City, CA, USA). The relative expression was calculated and normalized using the 2^−ΔΔCt^ method relative to 18S and/or GUSB.

### miRNA stability tests

Cells were transfected with ACTB UTR WT/MUT for 24h. Then cells were treated with Actinomycin-D for 4, 8, 12, 16, 20, 24 hours and total RNA was extracted with the miRNeasy Mini (Qiagen). The expression was detected by qRT-PCR.

### Plasmids Construction and luciferase reporter assay

A full-length fragment of the ACTB 3′UTR wild type or mutant was amplified. The PCR product was cloned into the pGL3-promoter luciferase-based plasmid (Promega) at the cloning site between KpnI and XhoI. The amplified fragment was verified by DNA sequencing. Hep3B cells were plated in 24-well plates and transfected with miR-1, miR-29a or NC together with ACTB WT or ACTB MUT. Cells were harvested after transfection 48h and the relative luciferase activity was measured by dual-luciferase reporter assay kit (Promega).

### Western blot

Lysates of individual cells (10 μg / lane) were separated by sodium dodecyl sulfate polyacrylamide gel (SDS-PAGE, 10 %) electrophoresis and transferred onto nitrocellulose membranes (Millipore Corporation, Billerica, MA). The membranes were then incubated in a SuperBlock T20 PBS Blocking Buffer (Thermo Fisher Scientific Inc., Waltham, MA). Subsequently, the membranes were incubated with primary antibodies (anti-GAPDH antibody, 1:1000, ab9485, and anti-beta-actin antibody, 1:1000, ab6276, abcam) according to the manufacturer’s instructions. Working solutions of the Pierce ECL Substrate and GE Healthcare ECL Reagent (Thermo Fisher Scientific) were prepared according to the manufacturers’ instructions. The membranes were digitally imaged using an enhanced chemiluminescence system.

### Cell migration/invasion assay

5×10^5^ cells were resuspended in culture medium without FBS and seeded in the upper chamber. Then the chamber was placed into a 24-well plate containing 600μl of culture media with 10% FBS. Approximately 48 h later, the cells were fixed with paraformaldehyde and stained with crystal violet, and the cells that passed through the membrane were counted.

### Xenograft study

All experimental procedures using animals were conducted in accordance with guidelines issued by the Animal Ethics Committee of Shanghai Eastern Hepatobiliary hospital (Shanghai, China). Hep3B cells stably expressing wildtype or mutant ACTB mRNA 3′UTR with fluorescent tag (5×10^6^ per injection) were injected into mice with introperitoneal injection.

### Statistical analysis

All statistical analyses were performed with GraphPad Prism software. The results (mean ±sd values) of all experiments were subjected to statistical analysis by *t* test. Values of *P* <0.05 were considered significant.

## Acknowledgements

We thank Dr. Mianshun Pan (Center of Radiation Oncology, Wujing Hospital, Shanghai 201103, China) for providing technical guidance. The project was supported by the grant from Science and Technology Commission of Shanghai Municipality (NO. 15411951900 to Feiling Feng).

## Author contributions

YL, H-BM and C-YS conducted the experiments. YL, F-LF and LY designed the experiments and wrote the paper. All authors revised, edited, and read the manuscript and approved the final manuscript.

## Conflict of interest

The authors declare that they have no conflict of interest.

